# Shotgun Proteomics of non-depleted and depleted hemolymph of human *Schistosomiasis snail vector; Biomphalaria glabrata*, revealed vital immune related proteins

**DOI:** 10.1101/2020.01.12.903708

**Authors:** Benson Otarigho, Mofolusho Falade, Castro-Borges William

## Abstract

**Background:** *Biomphalaria glabrata* is an important host in the transmission of human schistosomiasis in the Caribbean and South America. There are serious setbacks in the current control measures, hence need to explore alternatives.

**Objective:** To unveil *Biomphalaria glabrata* hemolymph proteome for possible detailed knowledge of its immunity in host-pathogen relationship.

**Methods:** In this study we performed a shotgun proteomic and bioinformatic analyses of the non-depleted and depleted [0.5 and 0.75% Trifluoroacetic acid (TFA) depletion] hemolymph of *B. glabrata* (LE strain).

**Results:** Our analysis was able to mine a total of 148 proteins from the hemolymph. The highest number of protein (148) was obtained from the 0.5% TFA-depleted sample, followed by 62 proteins from the 0.75% TFA-depleted sample and only 59 from non-depleted hemolymph.

**Conclusion:** Most of these proteins are novel and some of them may perform an important role in the immunity of this snail against invading pathogens.

## Introduction

*Biomphalaria glabrata* is an important host in the transmission of human schistosomiasis (1-7), which has been reported in 78 countries, amounting to more than 240 million people infected worldwide (8, 9). Although control measures involving the combined use of molluscicides and mass chemotherapy have been in place, the disease burden is still high and spreading especially in the developing countries (10-12). The absence of an ideal vaccine is partly due to lack of detailed understanding of the parasite/host interaction at both the human and snail stages, making reduction of schistosomiasis transmission around the world a difficult task. The knowledge on proteome composition of *B. glabrata* hemolymph, in which the internal defense mechanisms are impended, could provide clues in vaccine development (14, 15). Proteomic analysis of *Biomphalaria glabrata* hemocytes have being show (7, 16) however, the proteome of the hemolymph is still unknow.

Currently, advanced proteomic approaches have made it possible to identify proteins with greater ease and sensitivity (17, 18). Nevertheless, the efficiency of this MS-based proteomic technique can be hindered when dealing with complex samples such as biofluids, in which a few numbers of constituents dominate the bulk of the protein. In particular, the protein profile of *B. glabrata* hemolymph is dominated mostly by different proteoforms of hemocyanin (19, 20). This represents a problem for the detection of minor constituents such as those possibly involved in snail immunity and/or resistance against invading pathogens.

The findings from this study provides novel information on snail biology and may be of interest for researchers that will intend to explore biomarkers of infection and snail resistance for a deeper understanding of the complex schistosome-snail interaction.

## Materials and Methods

### Snails and hemolymph preparation

*B. glabrata* snails of the LE albino strain were used in this study. These snails were maintained in continuous culture in 40-liter aquarium which receive constant filtration and aeration at a constant temperature of 25°C with a 12: 12-hr 1ight:dark cycle and were fed with fresh leaf lettuce *ad libitum*. Hemolymph was obtained from healthy *B. glabrata* snails (12–15 mm shell diameter) by the head-foot retraction method [21, 22] and immediately placed on ice. Upon collection, hemolymph of each strain was dispensed into 1.5 mL microcentrifuge tubes containing cold, sterile Phosphate buffered saline (PBS) creating 1:1 dilution of PBS : hemolymph. 50ml of each sample was collected and check for the presence and absence of hemocytes before and after filtration respectively. The filtration was carried by passing each sample through 0.22um hydrophilic cellulose acetate filter membrane (EMD Millipore) by means of gentle press applied by an attached 5ml syringe. The filtrate was collected in 1.5ml microtube followed by addition of protease inhibitors (Protease Inhibitor Cocktail SetIII, EDTA-free; Calbiochem, San Diego, CA) to protect proteins from endogenous protease activities.

### Cytospinning, hemocyte fixation and staining

Cytocentrifuge was prepared with a labelled slide from the filtered hemolymph, chambered and blotted for each sample to be examined. 30ul of each sample was collected and diluted 1:1 in PBS and this was added to the slide chamber and spinned at 1500 rpm for 10 mins. Slides were carefully removed and allow to air dry prior to staining.

### Depletion of most abundant proteins

To deplete hemocynin and other most abundant proteins from the hemolymph. 5.3ul of prepared 10% trifluoroacetic acid (TFA) was added to 100ul of the hemolymph to have 0.5% depletion of most abundant protein while 8.1 ul of prepared 10% trifluoroacetic acid (TFA) was added to 100ul of the hemolymph to have 0.75% depletion of most abundant protein. This was followed by a gentle vortex and centrifuge at centrifuge at 2000g for 30mins at 4°C. The supernatant was collected and discard the pellet.

### 1D SDS-PAGE

To evaluate and visualize the effect of TFA depletion of the most abundant proteins from this snail hemolymph, 1D SDS-PAGE were carried out on 12% gel at constant 20 mA in a Miniprotean 2D-chamber (BioRad). Supernatants obtained after depletion were mixed in a ratio of 1: 1 with loading buffer and under reducing conditions were heated at 95°C for 5 min prior to loading onto the gels. Briefly, 1st lane was loaded with 3.76 mg/ml of non-depleted hemolymph, 2^nd^ lane was loaded with 2.32, mg/ml of 0.75% TFA depleted hemolymph and 3^rd^ lane was loaded with 2.03 mg/ml of 0.5% TFA depleted hemolymph. To facilitate comparisons, all samples were run simultaneously with low molecular weight markers (BioRad). After electrophoretic separation, the silver stained [23]. The imaged of the gel was scanned on Image Scanner III (GE Healthcare) and exported using the labscan tools version 6.01. This analysis intended the evaluation of (1) the number of bands in each depletion compared to the non-depleted hemolymph and (2) to compare the huge bands at the beginning of the gel.

### In-solution digestion and Nanoflow UHPLC system in peptide/protein identification

A total of 40ug of protein from each of the samples was digested with 1.6ug of sequencing grade trypsin following standard method (). All samples were analyzed using liquid chromatography electrospray ionization tandem mass spectrometry (LC-ESI-MS/MS) on a Ultimate^™^ 3000 series (Thermo Scientific DionexTM) coupled to a Q-Exactive mass spectrometer (Thermo Fisher Scientific, Bremen, Germany). The Q Exactive MS interfaced with nanoESI ionization as employed for MS analysis.

### *B. glabrata proteome* and annotation

The publicly available Proteome of *Biomphalaria glabrata* (BB02) version BglaB1.2 made available on 22th August, 2014, containing 14,137 protein sequences was downloaded from VectorBase, http://www.vectorbase.org, *Biomphalaria glabrata*, vector, BglaB1.2. and converted to fasta format using geneious version R8 (24). These sequences were functionally annotated on Blast2GO version 3.0 platform (25-28).

### Database searching and data analysis

RawMeat software version 2.0 and Thermo Xcalibur 2.2 Qual Browser were employed to assess the quality of the chromatograms of each Thermo Xcalibur .raw file and the efficiency of the protein digested with trypsin. All raw files were analyzed qualitatively and quantitatively using the Thermo Proteome Discoverer software platform version 1.4. The annotated proteome *B. glabrata* faste file was added to the platform fasta database and a label-free quantification with area detector and annotation workflow was used for the whole analysis. The analyzed results were filtered and exported in excel workbook (xlsx format) where the enrichment graphs on scatter plot were plotted after proper filtration. The number of proteins in each depleted and non-depleted samples was identified on the server http://bioinformatics.psb.ugent.be/webtools/Venn/ and the Venn diagram was generated on BioVenn (http://www.cmbi.ru.nl/cdd/biovenn/).

The annotations generated by Blast2go on the identified proteins were manually confirmed on Gene Ontology database (http://www. http://amigo.geneontology.org/amigo/search/ontology). These protein annotations were plotted on a pie chart using the excel software.

## Results

The head-foot retraction method used in the hemolymph collection from these snails yielded about 500μl of hemolymph from snails. We observed that the slide centrifuged method employed was able to remove the hemocytes and other cells that may be present from the hemolymph completely (Figure 1). This slide centrifuge method was to make sure that there were no cells or cell lysates in the samples before proceeding to depletion steps. The stereo microscopic images in Figure 1 (i) shows the present of hemocytes before the samples were slide centrifuged while Figure 1 (ii) shows the absence of hemocytes after samples were slide centrifuged. We observed different stages of development of the hemocyte (prohemocyte and oenocytoid) in the hemolymph of *B. glabrata* as presented in Figure 1 (i).

**Figure 1:**
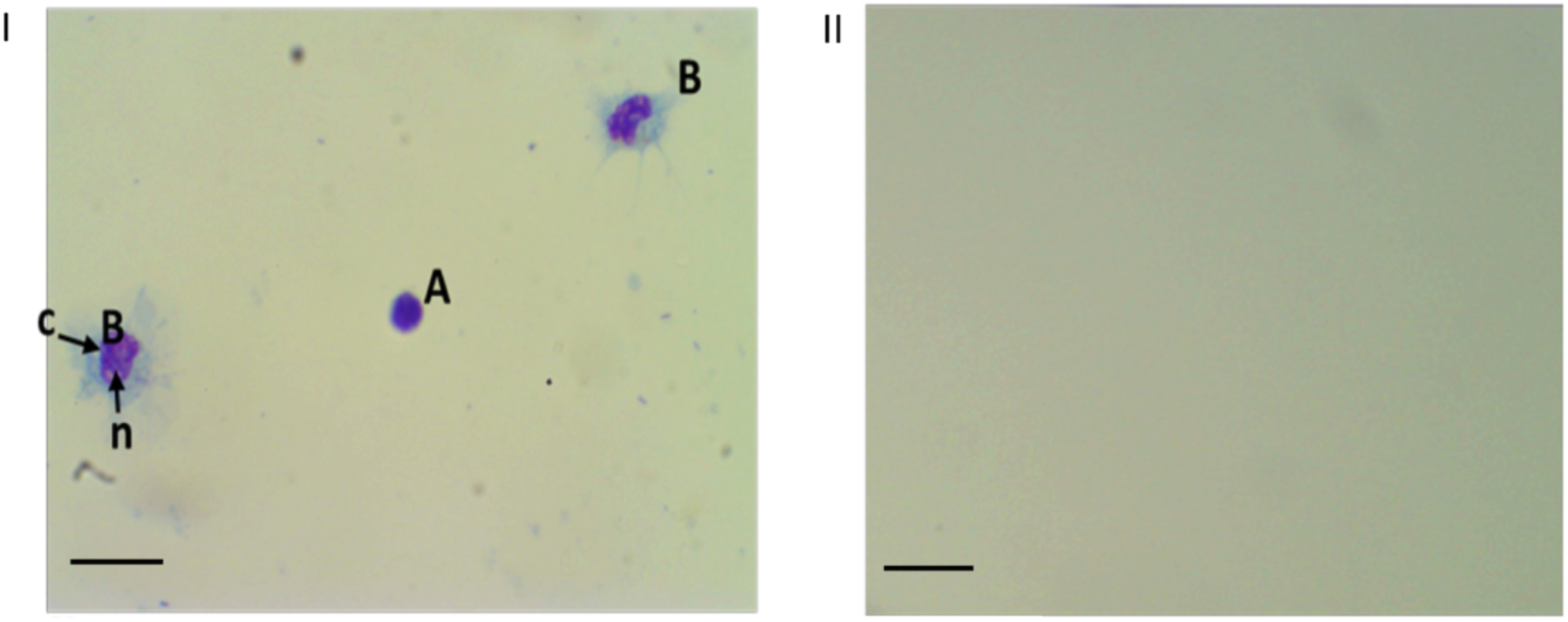
Stereo microscope images showing the present or absence of hemocytes before and after filtration. I show present of hemocytes before filtration while II. Shows the absence of hemocyte after filtration. (A), prohemocyte (B), oenocytoid. c = cytoplasm, n = nucleus. Bar = 100μm

The silver-stained 1D SDS PAGE gel in Figure 2 shows the comparative analysis of non-depleted and TFA-depleted hemolymph. The gel image revealed a significant enrichment for low molecular mass proteins (Mr < 116 kDa). In the non-depleted sample, there is a major huge dark band (Mr > 116 kDa) and a very few and faint bands are notice below 116kDa. In the depleted samples all bands show reproducible pattern. We noticed that that there was retained band in the stacking gel in the non-depleted sample, while it was not notice in the depleted samples. Again, among the two depleted samples, in the 0.5% TFA depleted sample, a band was noticed immediately the sample enter the separating gel but is absent in the 0.75% TFA depleted sample. However, the huge dark bands that are presented in non-depleted sample were highly reduced in depleted samples. Besides, there are much more proteins with lower molecular mass (Mr< 116 KDa) in the depleted samples when compare to non-depleted samples.

**Figure 2:**
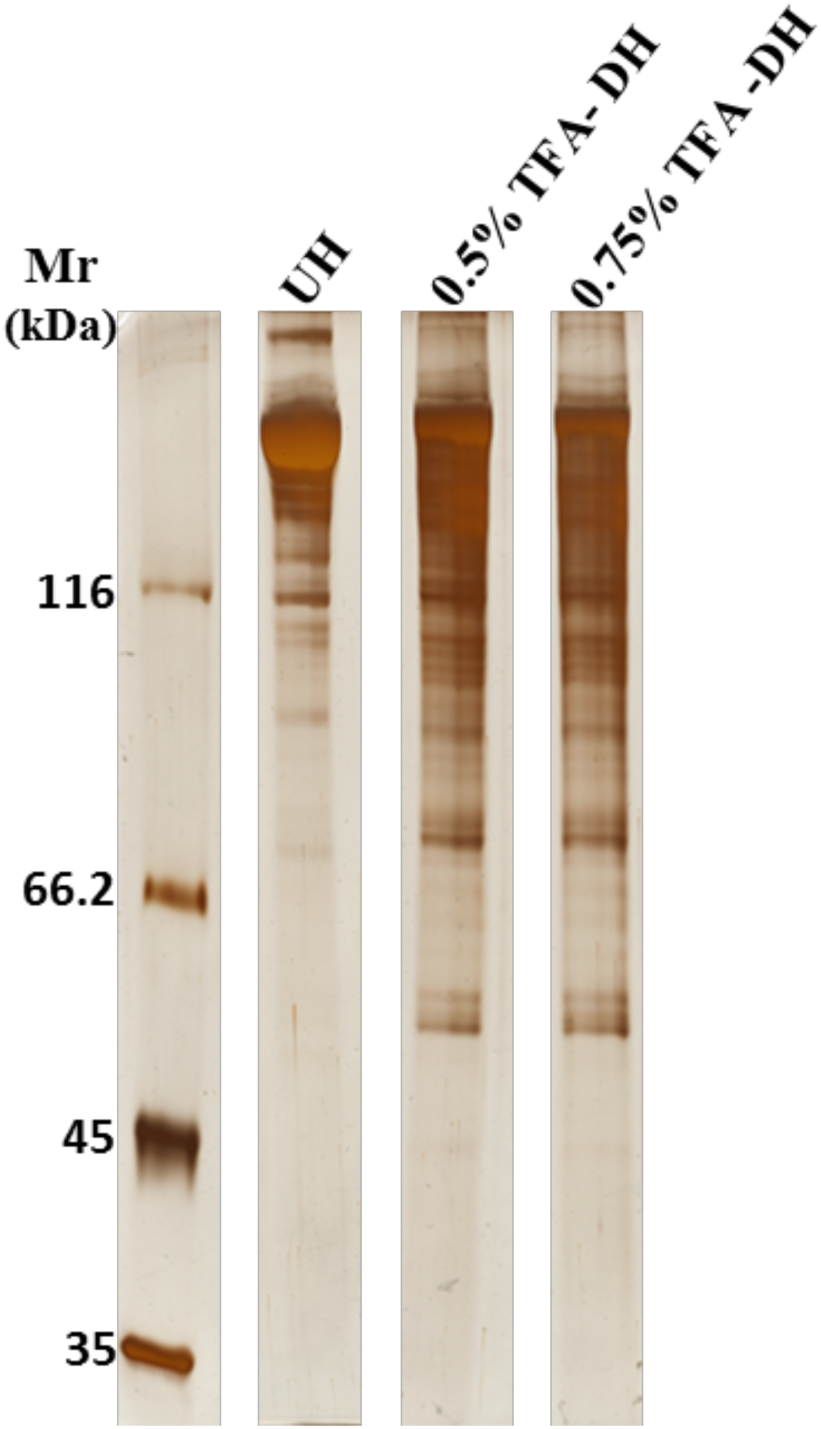
A comparative 1D gel analysis of non-depleted and TFA-depleted hemolymph revealed a significant enrichment for low molecular mass proteins (Mr < 116 kDa). 2μl of non-depleted hemolymph sample were loaded into 1st lane. 2.9μl of 0.5% TFA depleted sample were loaded into 2nd lane. Finally, 3.0μl of 0.75% TFA depleted sample were loaded into 3rd lane. *UH means non-depleted hemolymph *DH means Depleted hemolymph

The result show that all the data follow non-parametric rule and the coefficient of variation across three replicates in non-depleted, 0.5% and 0.75% TFA depletion were 15.2%, 26.13 and 24.81% respectively. The lowest variation across replication was observed in the non-depleted sample. Upon interrogation of the LC-MS/MS results with a specific *B. glabrata* proteome dataset on proteome discoverer 1.4 platform, a total of 148 proteins groups were identified across the three samples, at a false discovery rate of 1-5%, when peptide confidence and protein per peptide were set at medium and 1 respectively. Since one of our primary aims is to have a comprehensive analysis of *B. glabrata* hemolymph proteome composition using shotgun approach, the number of identified proteins in each and shared by samples are represented by a Venn diagram in Figure 3A. A great number of proteins (95) was obtained from the 0.5% TFA-depleted sample, whilst 0.75% TFA-depleted and non-depleted hemolymph resulted in 61 and 58 identities, respectively. The unique number of proteins in 0.5% TFA depleted, 0.75% TFA depleted and non-depleted sample are 50, 18 and 37 respectively, while the unique number of proteins shared between 0.5% and non-depleted, 0.75% TFA depleted and non-depleted, 0.5% TFA and 0.75% TFA depleted sample are 5, 3 and 27 respectively. A total of 13 proteins where shared by the three samples. We constructed a heat map chart to quantitatively demonstrate the effect of TFA depletion on proteins that are common across the three and between the two depleted samples (Figure 4). It is clear that different proteoforms of hemocyanins, which are the most abundant proteins were reduced in the depleted compared to non-depleted sample.

**Figure 3:**
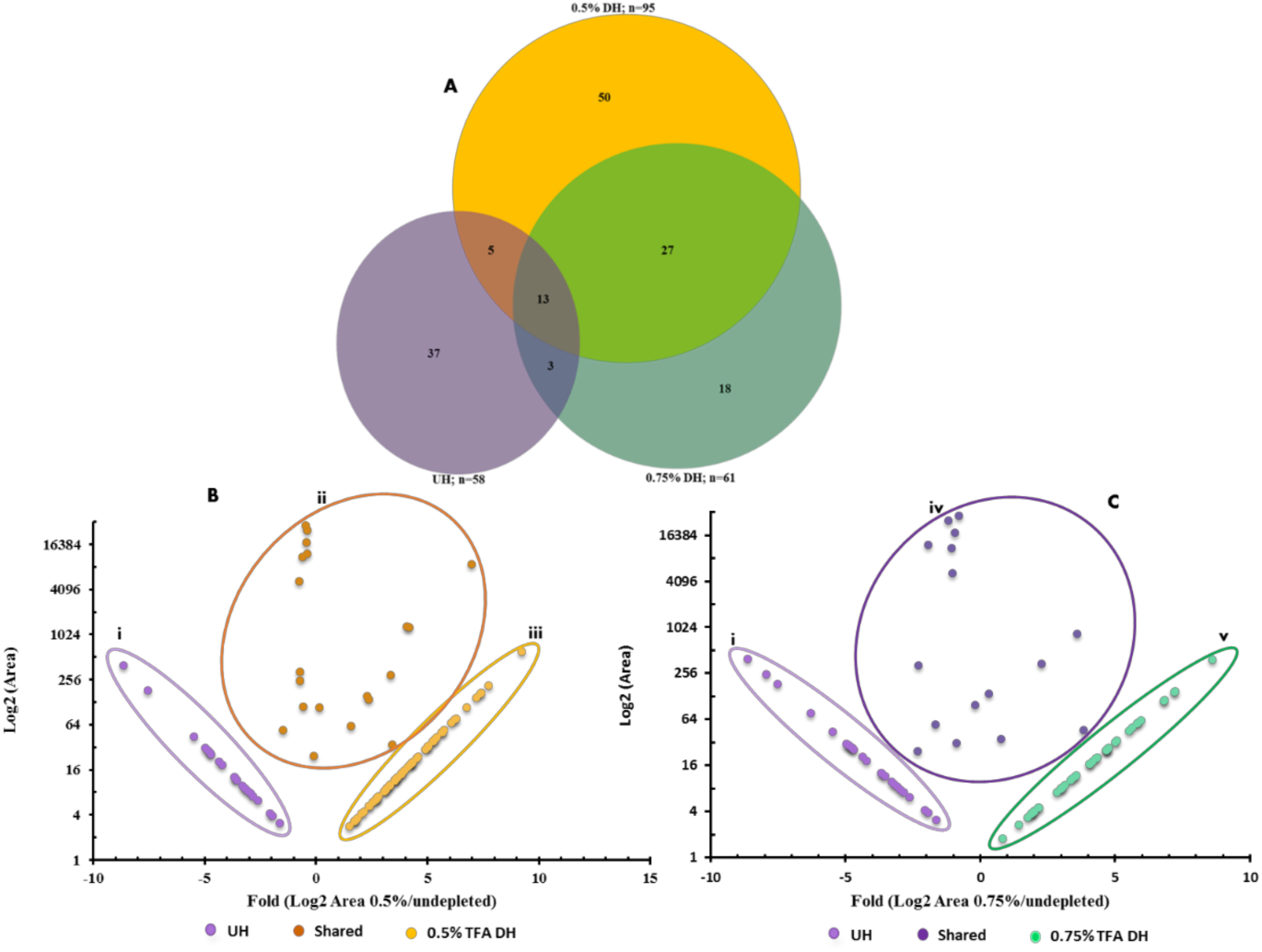
Venn diagram and scatter plot showing the number of proteins identified. (A) Total of 148 proteins was retrieved. A great number of proteins (95) was obtained from the 0.5% TFA-DH, whilst 0.75% TFA-DH and UH resulted in 61 and 58 identities, respectively. (B) Scatter plot quantitively displaying the identified proteins 0.5% TFA-DH against UH sample, proteins in group i and iii are exclusive to UH and 0.5% DH, while group ii were identified in both samples. (C) Scatter plot quantitively displaying the identified proteins 0.75% TFA DH against UH sample, proteins in group i and v are exclusive to UH and 0.75% DH, while group iv were identified in both samples. Significant proteins for B and C (Fold > 2). *UH means non-depleted hemolymph *DH means Depleted hemolymh

**Figure 4:**
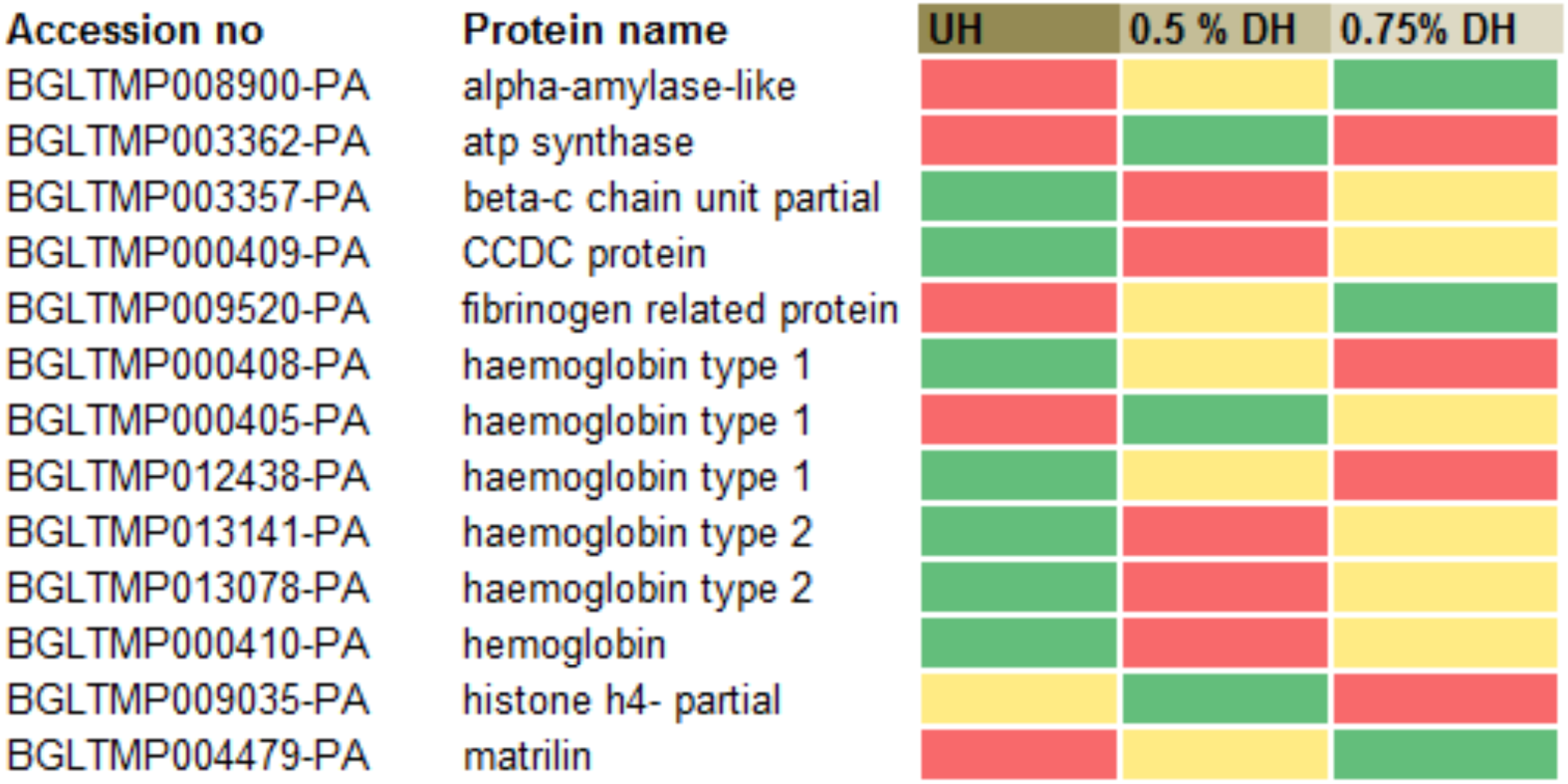
Heat map chart showing the quantitative effect of the depletion of 13 exclusive proteins in the three samples. The green, red and yellow cell show high, medium and low abundance respectively. *CCDC means coiled-coil domain-containing protein

To have a meaningful interpretation of the identified proteins in terms of abundance and enrichment, we employed scatter plot (Figure 3B & C). From this scatter plot supported by the Venn diagram in Figure 3A. We noticed that 0.5% TFA depletion was the better compare to 0.75% TFA depletion, since highest numbers of protein groups were identified. Most of these proteins were not identified in the non-depleted and 0.75% TFA depletion samples. From Figure 3A, the identified proteins in group (i) were only in non-depleted sample and the protein group in (iii) were identified in 0.5% TFA depleted sample only. Proteins in group (ii) were identified in both non-depleted and 0.5% TFA depleted sample, however their abundant is higher or lower either in non-depleted hemolymph or 0.75% TFA depleted sample. The comparative list of proteins identified in both 0.5% TFA depleted and non-depleted samples. From Figure 3B, the identified proteins in group (i) were only non-depleted sample and the protein in group (v) were identified in 0.75% TFA depleted sample only. Proteins in group (iv) were identified in both non-depleted and 0.75% TFA depleted sample, however there are some that their abundant is higher or lower either in non-depleted hemolymph or 0.75% TFA depleted sample. The comparative list of proteins identified in both 0.75% TFA depleted and non-depleted samples. Although, hemocyanin and it proteoforms were identified in all the three samples, their abundance in the depleted samples were much smaller in comparison to non-depleted samples.

For us to have an idea of the possible activities of the identified proteins, we classified them bases on their molecular functions within gene ontology categories, which are represented Figure 5. The majority (29%) of the identified proteins are enzymes, 11% are involve in transcription, 9% are involve in stress response/redox activities, 9% are involve in signaling, 7% are involve in defense against pathogens, 4% are involve in transport and 1% are involve in metabolism. Functions other than the mentions ones were classified as other processes and took 13%. The protein without annotation were classified as unknown function.

**Figure 5:**
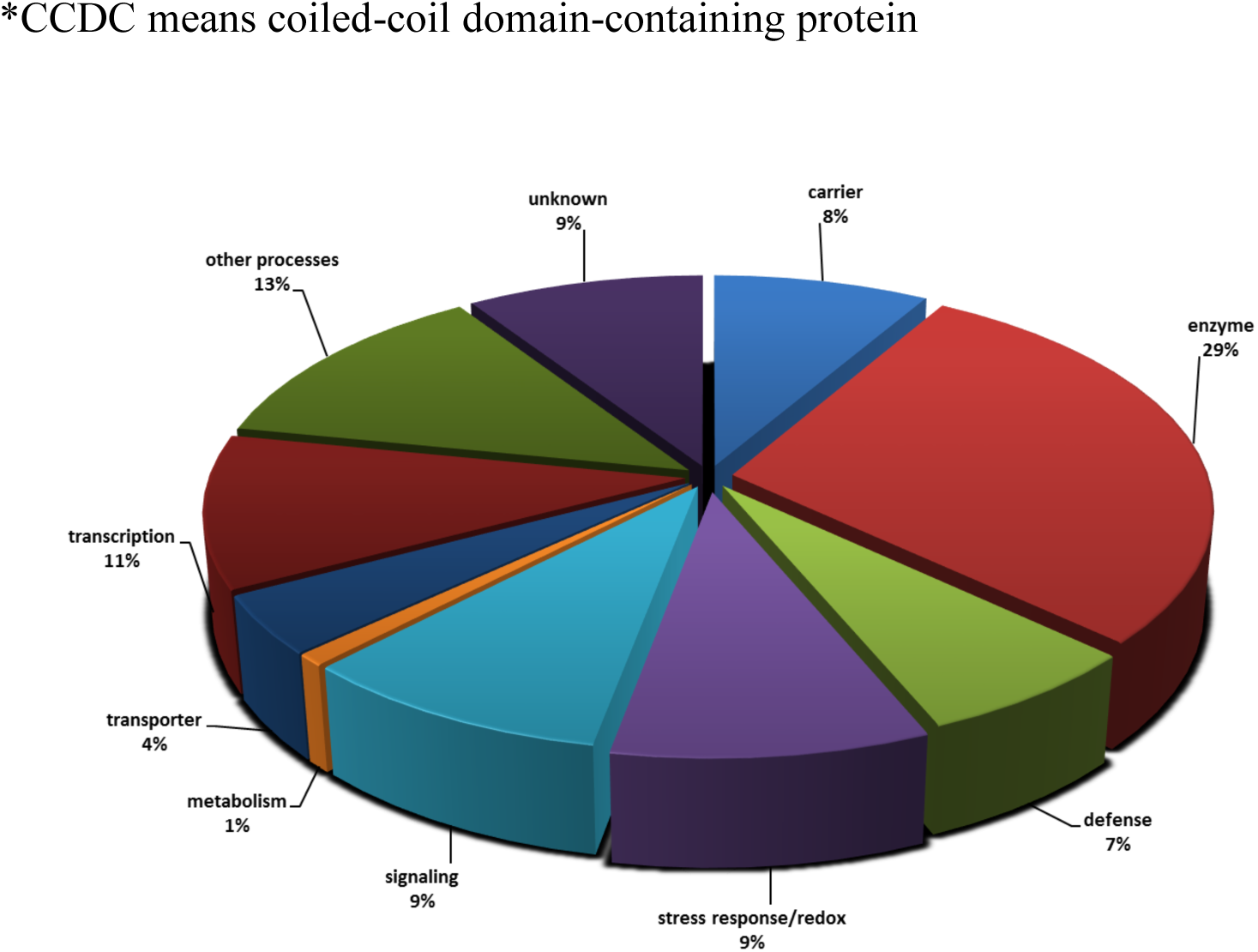
Pie-chart analyses of *B. glabrata* hemolymph proteome based on 10-gene ontology categories according to the number of constituents assigned to each category. The number of proteins used in this analysis was 148.

## Discussion

In general, biofluid and other tissues pose great challenge in the field of proteomics, due to masking of low abundance proteins by the high abundance ones [29-31]. In mammalian plasma and serum samples, albumin and IGs are the mostly abundant proteins [29-32] creating a great hinderance to detection other proteins. In the case of Mollusca the most common and abundant biofluid is the hemolymph and mucus. Studied had shown that the *B. glabrata* hemolymph is very important since it house the circulating hemocytes and protein components which play an immunological role in the relationship with parasites such, as *S. mansoni* developing larval, at the cellular and molecular level [33-35]. These circulating hemocytes and protein components dictate the susceptibility, resistivity of the snail against parasite invasion and survival [36-38]. However, identification of these proteins which are low in abundance from this snail hemolymph is a big challenge due to the high abundance of proteoforms of hemocyanin [38]. Besides, this will pose a persistent barrier in search for biomarkers of pathogen infection and host resistant.

In this study, we applied TFA depletion method to deplete the most abundant proteins in order to identified low abundant protein, then we unveiled the *B. glabrata* hemolymph proteome and each protein abundance. We noticed that the present of hemocytes in the samples will be a source of contaminant, so we modified known methods to get rid of the hemocytes. The foot-head retraction method used in collected of hemolymph was efficient and reduced cell damages, tissue debris or other contaminants. The slide centrifuge method used, completely remove hemocytes from *B. glabrata* hemolymph and this have not been used in molluscan research before now. This method was modified from [39, 40] where they isolate hemocytes from insects’ hemolymph such as mosquitoes and wild caterpillars. Majority of the hemocytes isolated from the hemolymph samples consisted of prohemocyte and oenocytoid. The hemocytes are very similar in shape to that of *Aedes aegypti* mosquitoes [39, 41].

In preliminary experiments, we tried many protocols, but all failed to effectively deplete hemocyanin from *B. glabrata* hemolymph. Fortunately, we found that TFA can effectively and selectively deplete hemocyanin from this snail hemolymph. We first optimized the depletion method regarding the amount of TFA, finally we found out that 0.5% and 0.75% TFA depletion was optimum for further analysis. When we increase the amount of TFA, so many proteins were depleted off, when the amount of TFA was lower 0.5%, it was not significantly different from the non-depleted sample. The proteins bands in the non-depleted samples in first three lanes of the 1D SDS PAGE gel (Figure 2) agree with that of [20]. The huge dark band is suggested to be proteoforms of hemocyanin, which are oxygen transporters in this snail. Because molluscan hemocyanin molecules are oligomers composed of many protein subunits [42], these form different bands that overlap forming this huge dark bands. The much more bands of low molecular weight seen in the depleted samples show the effect of the TFA depletion method. This depletion method seems to be more efficient when compare to when ultrafiltration centrifuge was used to reduce the abundance of hemocyanin [7, 20] based on the many bands seen in silver stained 1D SDS PAGE gel (Figure 2).

## Conclusion

A number of novel proteins were identified from the hemolymph of this schistosomiasis snail vector and most of them were proposed to be crucially involve involved in immunity and others with unknown function were identified. Our investigation opens new avenues for elucidation of the schistosome-snail interaction. Besides, based on the findings in this work and apply this shot proteomic workflow, the search for biomarkers of *S. mansoni* infection in this snail hemolymph is on the way.

## Supporting information

Supplementary Table 1_05 depletion

Supplementary Table 2_075 depletion

Supplementary Table 3 whole

## Acknowledgements

The authors would like to acknowledge the financial support provided by Conselho Nacional de Desenvolvimento Científico e Tecnológico (CNPq) and The World Academy of Sciences (TWAS) with the award number: 3240274297.

## References

1. Pimentel, D. Life history of Australorbis glabratus, the intermediate snail host of Schistosoma mansoni in Puerto Rico. Ecology 38, 576–580 (1957).

2. Steffey, E. P. & Howland, J. D. Isoflurane potency in the dog and cat. Am. J. Vet. Res. 38, 1833–1836 (1977).

3. Pointier, J. P. The introduction of Melanoides tuberculata (Mollusca: Thiaridae) to the island of Saint Lucia (West Indies) and its role in the decline of Biomphalaria glabrata, the snail intermediate host of Schistosoma mansoni. Acta Trop. 54, 13–18 (1993).

4. Raccurt, C. P., Sodeman Jr, W. A., Rodrick, G. L. & Boyd, W. P. Biomphalaria glabrata in Haiti. Trans. R. Soc. Trop. Med. Hyg. 79, 455–457 (1985).

5. Sturrock, R. F. Ecological notes on habitats of the freshwater snail Biomphalaria glabrata, intermediate hosts of Schistosoma mansoni on St. Lucia, West Indies. Carib J Sci 14, 149–162 (1974).

6. Wethington, A. R. et al. Population genetic structure of Biomphalaria glabrata in a schistosomiasis-endemic region in Brazil. J. Molluscan Stud. 73, 45–52 (2007).

7. Dinguirard, N. et al. Proteomic Analysis of Biomphalaria glabrata Hemocytes During in vitro Encapsulation of Schistosoma mansoni Sporocysts. Front. Immunol. 9, 2773 (2018).

8. El Ridi, R. & Tallima, H. Vaccine-induced protection against murine schistosomiasis mansoni with larval excretory-secretory antigens and papain or type-2 cytokines. J. Parasitol. 99, 194–202 (2013).

9. Organization, W. H. Schistosomiasis: number of people receiving preventive chemotherapy in 2012. Wkly. Epidemiol. Rec. Relev. épidémiologique Hebd. 89, 21–28 (2014).

10. Sturrock, R. F. Schistosomiasis epidemiology and control: how did we get here and where should we go? Mem. Inst. Oswaldo Cruz 96, 17–27 (2001).

11. Angelo, T. et al. Population Abundance and Disease Transmission Potential of Snail intermediate hosts of Human Schistosomiasis in Fishing Communities of Mwanza Region, North-western, Tanzania. Int. J. Sci. Res. 3, 1230–1236 (2014).

12. Chala, B. & Torben, W. An epidemiological Trend of Urogenital Schistosomiasis in ethiopia. Front. public Heal. 6, 60 (2018).

13. Bergquist, N. R. & Colley, D. G. Schistosomiasis vaccine: research to development. Parasitol. Today 14, 99–104 (1998).

14. Mager, W. H. & Ferreira, P. M. Stress response of yeast. Biochem. J. 290, 1 (1993).

15. Barbosa, L., Silva, L. M., Coelho, P. M. Z., Santos, S. R. & Fortes-Dias, C. L. Primary culture of the region of the amebocyte-producing organ of the snail Biomphalaria glabrata, the intermediate host of Schistosoma mansoni. Mem. Inst. Oswaldo Cruz 101, 639–643 (2006).

16. Adema, C. M., Hertel, L. A. & Loker, E. S. Evidence From Two Planorbid Snails.pdf. Parasitology (1999).

17. Nilsson, T. et al. Mass spectrometry in high-throughput proteomics: ready for the big time. Nat. Methods 7, 681 (2010).

18. Pandeswari, P. B. & Sabareesh, V. Middle-down approach: a choice to sequence and characterize proteins/proteomes by mass spectrometry. RSC Adv. 9, 313–344 (2019).

19. Lieb, B. et al. Red blood with blue-blood ancestry: intriguing structure of a snail hemoglobin. Proc. Natl. Acad. Sci. 103, 12011–12016 (2006).

20. Yoshino, T. P. et al. Glycotope sharing between snail hemolymph and larval schistosomes: larval transformation products alter shared glycan patterns of plasma proteins. PLoS Negl. Trop. Dis. 6, e1569 (2012).

21. Sminia, T. & Barendsen, L. A comparative morphological and enzyme histochemical study on blood cells of the freshwater snails Lymnaea stagnalis, Biomphalaria glabrata, and Bulinus truncatus. J. Morphol. 165, 31–39 (1980).

22. Sminia, T. Structure and function of blood and connective tissue cells of the fresh water pulmonate Lymnaea stagnalis studied by electron microscopy and enzyme histochemistry. Zeitschrift für Zellforsch. und mikroskopische Anat. 130, 497–526 (1972).

23. Shevchenko, A., Wilm, M., Vorm, O. & Mann, M. Mass spectrometric sequencing of proteins from silver-stained polyacrylamide gels. Anal. Chem. 68, 850–858 (1996).

24. Kearse, M. et al. Geneious Basic: an integrated and extendable desktop software platform for the organization and analysis of sequence data. Bioinformatics 28, 1647–1649 (2012).

25. Conesa, A. et al. Blast2GO: a universal tool for annotation, visualization and analysis in functional genomics research. Bioinformatics 21, 3674–3676 (2005).

26. Conesa, A. & Götz, S. Blast2GO: A comprehensive suite for functional analysis in plant genomics. Int. J. Plant Genomics 2008, (2008).

27. Götz, S. et al. B2G-FAR, a species-centered GO annotation repository. Bioinformatics 27, 919–924 (2011).

28. Götz, S. et al. High-throughput functional annotation and data mining with the Blast2GO suite. Nucleic Acids Res. 36, 3420–3435 (2008).

29. Schenk, S., Schoenhals, G. J., de Souza, G. & Mann, M. A high confidence, manually validated human blood plasma protein reference set. BMC Med. Genomics 1, 41 (2008).

30. Sharma, V., Jaishankar, A., Wang, Y.-C. & McKinley, G. H. Rheology of globular proteins: apparent yield stress, high shear rate viscosity and interfacial viscoelasticity of bovine serum albumin solutions. Soft Matter 7, 5150–5160 (2011).

31. Carlyle, B., Trombetta, B. & Arnold, S. Proteomic Approaches for the Discovery of Biofluid Biomarkers of Neurodegenerative Dementias. Proteomes 6, 32 (2018).

32. Anderson, N. L. et al. The human plasma proteome: a nonredundant list developed by combination of four separate sources. Mol. Cell. Proteomics 3, 311–326 (2004).

33. Yoshino, T. P. & Vasta, G. R. Parasite-invertebrate host immune interactions. In Invertebrate Immune Responses 125–167 (Springer, 1996).

34. Horák, P. & Knaap, W. P. W. van der. Lectins in snail-trematode immune interactions: a review. Folia Parasitol. (Praha). 44, 161–172 (1997).

35. Sturrock, R. F. The schistosomes and their intermediate hosts. Schistosomiasis 3, 7–83 (2001).

36. Bayne, C. J., Buckley, P. M. & DeWan, P. C. Schistosoma mansoni: cytotoxicity of hemocytes from susceptible snail hosts for sporocysts in plasma from resistant Biomphalaria glabrata. Exp. Parasitol. 50, 409–416 (1980).

37. Bayne, C. J., Buckley, P. M. & DeWan, P. C. Macrophagelike hemocytes of resistant Biomphalaria glabrata are cytotoxic for sporocysts of Schistosoma mansoni in vitro. J. Parasitol. 413–419 (1980).

38. Yoshino, T. P., Wu, X.-J., Gonzalez, L. A. & Hokke, C. H. Circulating Biomphalaria glabrata hemocyte subpopulations possess shared schistosome glycans and receptors capable of binding larval glycoconjugates. Exp. Parasitol. 133, 28–36 (2013).

39. Qayum, A. A. & Telang, A. A protocol for collecting and staining hemocytes from the yellow fever mosquito Aedes aegypti. J. Vis. Exp. JoVE (2011).

40. Stoepler, T. M., Castillo, J. C., Lill, J. T. & Eleftherianos, I. A simple protocol for extracting hemocytes from wild caterpillars. J. Vis. Exp. JoVE (2012).

41. Hillyer, J. F. & Christensen, B. M. Characterization of hemocytes from the yellow fever mosquito, Aedes aegypti. Histochem. Cell Biol. 117, 431–440 (2002).

42. Markl, J. Evolution of molluscan hemocyanin structures. Biochim. Biophys. Acta (BBA)-Proteins Proteomics 1834, 1840–1852 (2013).

43. Hanington, P. C. & Zhang, S.-M. The primary role of fibrinogen-related proteins in invertebrates is defense, not coagulation. J. Innate Immun. 3, 17–27 (2011).

